# Quercetin alters redox homeostasis, induces MAPK activation and promotes DNA damage in vGPCR-transformed endothelial cells

**DOI:** 10.64898/2026.09.30.755570

**Authors:** Gabriel Principe, Athina Maniscalchi, Silvina Tiburzi, Virginia Lezcano, Gabriela A. Salvador, Verónica González-Pardo

**Affiliations:** Departamento de Biología, Bioquímica y Farmacia, Universidad Nacional del Sur (UNS), San Juan 670, 8000, Bahía Blanca, Argentina; Instituto de Ciencias Biológicas y Biomédicas del Sur (INBIOSUR), UNS-Consejo Nacional de Investigaciones Científicas y Técnicas (CONICET), San Juan 671, 8000, Bahía Blanca, Argentina; Instituto de Investigaciones Bioquímicas de Bahía Blanca (INIBIBB), UNS-CONICET, Camino La Carrindanga km 7, 8000, Bahía Blanca, Argentina

**Keywords:** Quercetin, Kaposi’s sarcoma, viral oncogenesis, oxidative stress, MAPK signaling, DNA damage

## Abstract

Kaposi’s sarcoma (KS) is an angioproliferative tumor associated with infection by human herpesvirus 8 (HHV-8), also known as Kaposi’s sarcoma-associated herpesvirus (KSHV). Among the viral genes involved in KS pathogenesis, the viral G protein-coupled receptor (vGPCR) plays a key role in oncogenic transformation and angiogenesis, and its sustained activity contributes to tumor development and maintenance. Quercetin (QUE), a naturally occurring flavonoid with well-recognized anticancer properties, has been shown to reduce cell viability and proliferation and to induce apoptosis in vGPCR-transformed endothelial cells. This study further investigated the mechanisms underlying the antineoplastic effects of QUE in vGPCR cells, an *in vitro* model of Kaposi’s sarcoma. Initially, QUE promoted the activation of MAPK signaling pathways in vGPCR cells, increasing the phosphorylation levels of ERK1/2, JNK1/2, and p38, without affecting AKT phosphorylation. In parallel, QUE increased intracellular oxidant levels, followed by lipid peroxidation and DNA damage, supporting a link between oxidative imbalance and genotoxicity. However, increased total glutathione content, and unaltered *Gpx4* mRNA expression, suggest the induction of oxidative stress and compensatory antioxidant responses. Furthermore, increased intracellular oxidant levels were associated with MAPK signaling activation, particularly JNK1/2, whereas ERK1/2 and p38 activation may reflect a compensatory response that is insufficient to overcome QUE-induced cytotoxicity. Taken together, these findings indicate that QUE exerts antiproliferative effects in vGPCR cells through a mechanism involving oxidative stress, MAPK activation, and DNA damage, supporting its potential relevance in the study of therapeutic strategies for virus-associated malignancies.

**Highlights:**

- QUE increased intracellular oxidant levels, lipid peroxidation, and total glutathione content.
- QUE activated MAPK signaling without affecting AKT phosphorylation.
- Pharmacological inhibition suggested a major role for JNK1/2 in oxidant accumulation.
- QUE promoted DNA damage in vGPCR-transformed endothelial cells.

## Introduction

In recent years, increasing attention has been directed toward naturally derived bioactive compounds because of their association with beneficial effects on human health and well-being (1). In this context, the term “nutraceuticals” refers to compounds with both nutritional and therapeutic potential, characterized by biological activity and safety for consumption (2). Among nutraceutical compounds, polyphenols represent one of the most abundant and diverse groups, comprising more than 40,000 plant secondary metabolites involved in defense responses against pathogens and environmental stress. In humans, polyphenols have been associated with broad therapeutic effects due to their diverse bioactivities (3). These compounds can modulate the activity of multiple enzymes and interfere with intracellular signaling pathways and other cellular processes (4). Moreover, because of their physicochemical properties, polyphenols can participate in cellular redox reactions, contributing to their beneficial antioxidant effects (5).

Quercetin (QUE), also known as 3,3′,4′,5,7-pentahydroxyflavone, is a polyphenolic flavonoid widely distributed in fruits and vegetables, including onions, apples, berries, and capers (6). QUE exhibits a wide range of biological activities and has been extensively studied in relation to several pathological conditions, including cancer (7), Alzheimer’s disease (8), and obesity (9).

Kaposi’s sarcoma (KS) is a virus-associated malignancy first described in 1872 by the Hungarian physician and dermatologist Moritz Kaposi (10). Its etiological agent, human herpesvirus 8 (HHV-8), also known as Kaposi’s sarcoma-associated herpesvirus (KSHV), was identified in 1994 (11), at a time when epidemiological evidence and the high incidence of KS in patients with AIDS strongly suggested the involvement of a sexually transmitted infectious agent distinct from HIV (12). A key viral gene expressed during the lytic replication phase encodes the viral G protein-coupled receptor (vGPCR), which contributes to oncogenesis and angiogenesis through the activation of multiple signaling pathways due to its constitutive activity (13–15). Indeed, tumor cells depend on vGPCR activity and on the survival of vGPCR-expressing cells, making this receptor essential for tumor transformation and maintenance (16,17). Therefore, vGPCR and its downstream signaling pathways represent relevant targets for the development of new therapeutic strategies against KS.

Although the antineoplastic activity of QUE has been widely investigated (18), and its antiviral activity has also been reported, mainly through inhibition of viral entry and replication (19), its impact on virus-associated cancers remains comparatively underexplored. In this context, our group has previously shown that QUE significantly reduced the proliferation and viability of vGPCR-transformed endothelial cells *in vitro* and inhibited tumor growth *in vivo*, supporting its antineoplastic activity in this model of viral oncogenesis (20). Hence, the present study aimed to further investigate the therapeutic potential of QUE in this model, with special attention to oxidative stress, MAPK signaling, and DNA damage as potential mechanisms underlying QUE-induced cytotoxicity. Thus, this work may contribute to the understanding and development of new strategies for the treatment of oncovirus-associated malignancies.

## Materials and Methods

### Chemicals and reagents

Cell culture was performed using Dulbecco’s modified Eagle’s medium (DMEM; Sigma-Aldrich, St. Louis, MO, USA) supplemented with fetal bovine serum (FBS; Natocor, Córdoba, Argentina). G418 antibiotic was purchased from Sigma-Aldrich (St. Louis, MO, USA). Quercetin (QUE) was obtained from MP Biomedicals (Solon, OH, USA). The fluorescent probe 2′,7′-dichlorofluorescein diacetate (DCFDA), agarose, low-melting-point agarose, and Ellman’s reagent [5,5′-dithiobis(2-nitrobenzoic acid); DTNB] were purchased from Sigma-Aldrich (St. Louis, MO, USA). BioZOL reagent was obtained from Thermo Fisher Scientific (Waltham, MA, USA). M-MLV reverse transcriptase was purchased from PB-L (Buenos Aires, Argentina). SsoAdvanced Universal SYBR Green Supermix was obtained from Bio-Rad Laboratories (Hercules, CA, USA). The kinase inhibitors SB203580, PD98059, SP600125, and LY294002 were obtained from Tocris Bioscience (Ellisville, MO, USA). Primary antibodies against phospho-ERK1/2, phospho-JNK, and phospho-AKT were purchased from Cell Signaling Technology (Danvers, MA, USA). Antibodies against phospho-p38 and α-Tubulin were obtained from Invitrogen (Thermo Fisher Scientific, Rockford, IL, USA). HRP-conjugated anti-rabbit secondary antibody was purchased from Santa Cruz Biotechnology (Santa Cruz, CA, USA). All other reagents were of analytical grade.

### Cell line and culture conditions

The experimental model of Kaposi’s sarcoma consisted of SV40-immortalized murine endothelial cells stably expressing the vGPCR full length receptor (vGPCR cells) as previously described (20). vGPCR cells were cultured in DMEM supplemented with 5% fetal bovine serum (FBS), 4.5 g/L glucose and selected with 500 μg/mL G418. Cells were allowed to grow in a metabolic incubator at 37 °C and 5% CO_2_ and the culture medium was replaced every two days.

### Trypan blue exclusion assay

vGPCR cells were seeded in 24-well plates at a density of 8 × 10^3^ cells/well. Following treatment with 30 µM QUE for 24 or 72 h, cells were trypsinized, stained with 0.04% trypan blue, and viable cells were counted using a Neubauer hemocytometer under a bright-field microscope.

### Determination of intracellular oxidants

Intracellular oxidant levels were determined using the fluorogenic probe 2′,7′-dichlorofluorescein diacetate (DCFDA). Cells were seeded in 48-well plates (3 × 10^3^ cells/well) and treated as indicated. H_2_O_2_ (0.5 mM, 45 min) was used as a positive control. Following incubation with 10 µM DCFDA for 30 min at 37 °C, fluorescence was measured using a Synergy HT microplate reader (BioTek; Ex/Em: 485/528 nm). Fluorescence values were normalized to total protein content.

### Lipid peroxidation assay

Lipid peroxidation was assessed by measuring thiobarbituric acid reactive substances (TBARS). Cells were seeded in 100-mm culture dishes (1.5 × 10^5^ cells/dish). Following the indicated treatments, cell lysates were mixed with 30% trichloroacetic acid, 5 N HCl, and 0.75% thiobarbituric acid. Samples were heated at 100 °C for 15 min, centrifuged to remove precipitated material, and the absorbance of the supernatant was measured spectrophotometrically at 535 nm. TBARS levels were normalized to total protein content (21).

### Determination of total glutathione (GSH)

GSH content was determined using a colorimetric assay based on the reaction of GSH with 5,5′-dithiobis(2-nitrobenzoic acid) (DTNB), which forms a yellow-colored chromophore. Cells were seeded in 100-mm culture dishes (1.5 × 10^5^ cells/dish). Following the indicated treatments, cell lysates were processed for the DTNB assay, and the absorbance was measured spectrophotometrically at 412 nm. GSH content was normalized to total protein content (21).

### Real-time quantitative PCR (RT-qPCR)

Total RNA was isolated using BioZOL reagent. Complementary DNA (cDNA) was synthesized from 1 μg of total RNA with M-MLV reverse transcriptase, following the manufacturer’s recommendations. RT-qPCR reactions were performed with SsoAdvanced Universal SYBR Green Supermix on a Rotor-Gene 6000 real-time PCR system (Corbett Research, Australia). The primer pairs used for the amplification of *Gpx4* and the reference gene *Gapdh* were as follows: *Gpx4* forward, 5′-TCTGTGTAAATGGGGACGATGC-3′, and reverse, 5′-TCTCTATCACCTGGGGCTCCTC-3′; *Gapdh* forward, 5′-TTCACCACCATGGAGAAGGC-3′, and reverse, 5′-AGTGATGGCATGGACTGTGGTC-3′. Relative mRNA expression was determined according to the 2^ΔΔCt^ method, employing *Gapdh* as the endogenous reference gene (22).

### SDS-PAGE and Western blot

Cells were lysed in ice-cold lysis buffer containing protease and phosphatase inhibitors. Protein concentration was determined using the Bradford assay (23). Equal amounts of protein (15–30 µg) were resolved by SDS-PAGE and transferred onto PVDF membranes. After blocking, membranes were incubated overnight at 4 °C with the indicated primary antibodies at the following dilutions: anti-pAKT (1:1,000), anti-pERK1/2 (1:1,000), anti-pJNK1/2 (1:250), anti-pp38 (1:1,000), anti-α-Tubulin (1:1,000); followed by HRP-conjugated anti-rabbit secondary antibody (1:5,000). Immunoreactive bands were detected using the ECL Western Blotting Detection Reagents (Amersham, GE Healthcare, UK) and quantified by densitometric analysis using ImageJ (24). When required, membranes were stripped and blotted with additional antibodies.

### Comet assay

DNA damage was evaluated using the alkaline comet assay. Cells were seeded in 35-mm dishes (3 × 10^4^ cells/dish). H_2_O_2_ (0.5 mM, 1 h) was used as a positive control. Following treatment, cells were processed according to the alkaline comet assay protocol described by Tiburzi and col. (25). Comets were visualized using an Olympus BX51 fluorescence microscope, and comet tail length was quantified using the OpenComet plugin for ImageJ (24,26,27).

### Statistical analysis

Data normality and homogeneity of variance were evaluated using the Shapiro–Wilk and Levene tests, respectively. Parametric data were analyzed using one-way ANOVA followed by Dunnett’s multiple-comparisons test or Student’s *t*-test, whereas nonparametric data were analyzed using the Kruskal–Wallis and Mann–Whitney U tests. Additionally, linear mixed-effects models were fitted to integrate data from independent experiments, and *post hoc* multiple-comparisons were performed using Tukey’s or Šídák’s adjustment, depending on the experimental design. All experiments were independently repeated at least three times. Statistical significance was indicated as follows: *p < 0.05; **p < 0.01; ***p < 0.001.

## Results

### QUE reduces vGPCR cell proliferation

First, based on the IC_50_ value previously determined for QUE in vGPCR cells after 48 h of treatment (20), the effect of QUE on cell proliferation was evaluated at shorter and longer exposure times. vGPCR cells were treated with QUE (30 µM) or vehicle (DMSO < 0.1%) for 24 or 72 h, and viable cells were quantified using the trypan blue exclusion assay. As shown in **Figure 1**, QUE treatment significantly reduced the number of viable cells compared with the control condition at both time points, indicating that this concentration exerts antiproliferative effects as early as 24 h of exposure.

**Figure 1.**
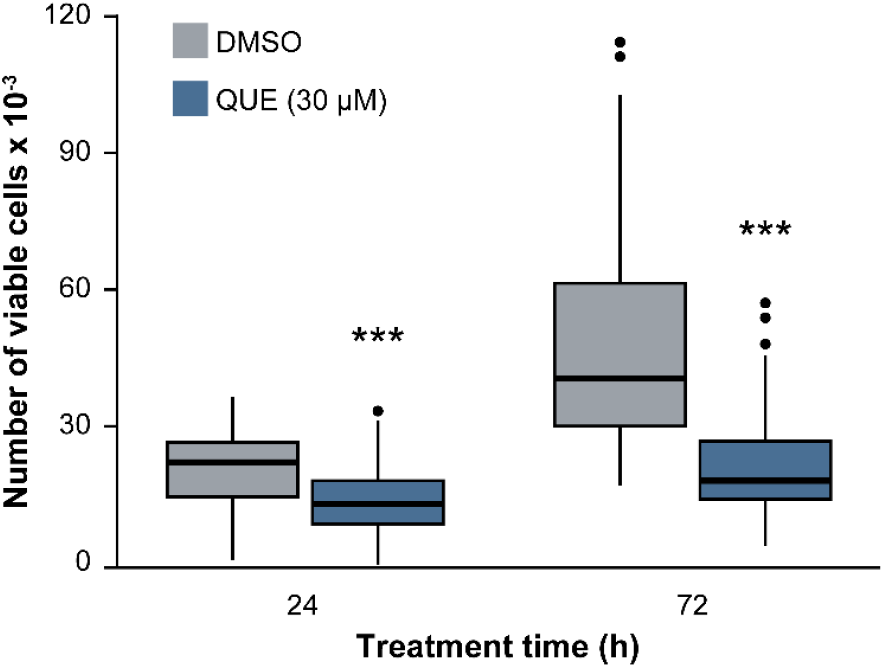
QUE effects on cell proliferation. vGPCR cells were treated with QUE (30 µM) or vehicle (DMSO < 0.1%) for 24 or 72 h, and cell proliferation was evaluated using the trypan blue exclusion assay. Data is presented as boxplot from three independent experiments. Statistical analysis was performed using a generalized linear mixed-effects model (***p < 0.001).

### QUE modulates ERK1/2, JNK1/2, and p38 activation in vGPCR cells

Mitogen-activated protein kinases (MAPKs) are involved in most signal transduction pathways regulating eukaryotic cell function, allowing coordinated cellular responses to a wide range of stimuli (28). Additionally, among the intracellular pathways activated by vGPCR, the PI3K/AKT signaling pathway represents one of the most important oncogenic mechanisms (29). Thus, to investigate the involvement of MAPKs (p38, JNK1/2, and ERK1/2) and AKT in QUE mechanism of action, vGPCR cells were treated with increasing concentrations of QUE (10–50 µM) or vehicle (DMSO < 0.1%) for 48 h. Thereafter, the protein levels of the phosphorylated forms of these kinases were assessed by Western blot as an indicator of their activation status. As shown in **Figure 2**, QUE treatment did not alter AKT phosphorylation at any concentration, while higher QUE concentrations (30 and 50 µM) significantly increased the phosphorylation levels of ERK1/2, JNK1/2, and p38. Notably, phosphorylated p38 (pp38) exhibited the most pronounced response, increasing by several orders of magnitude relative to the control condition.

**Figure 2.**
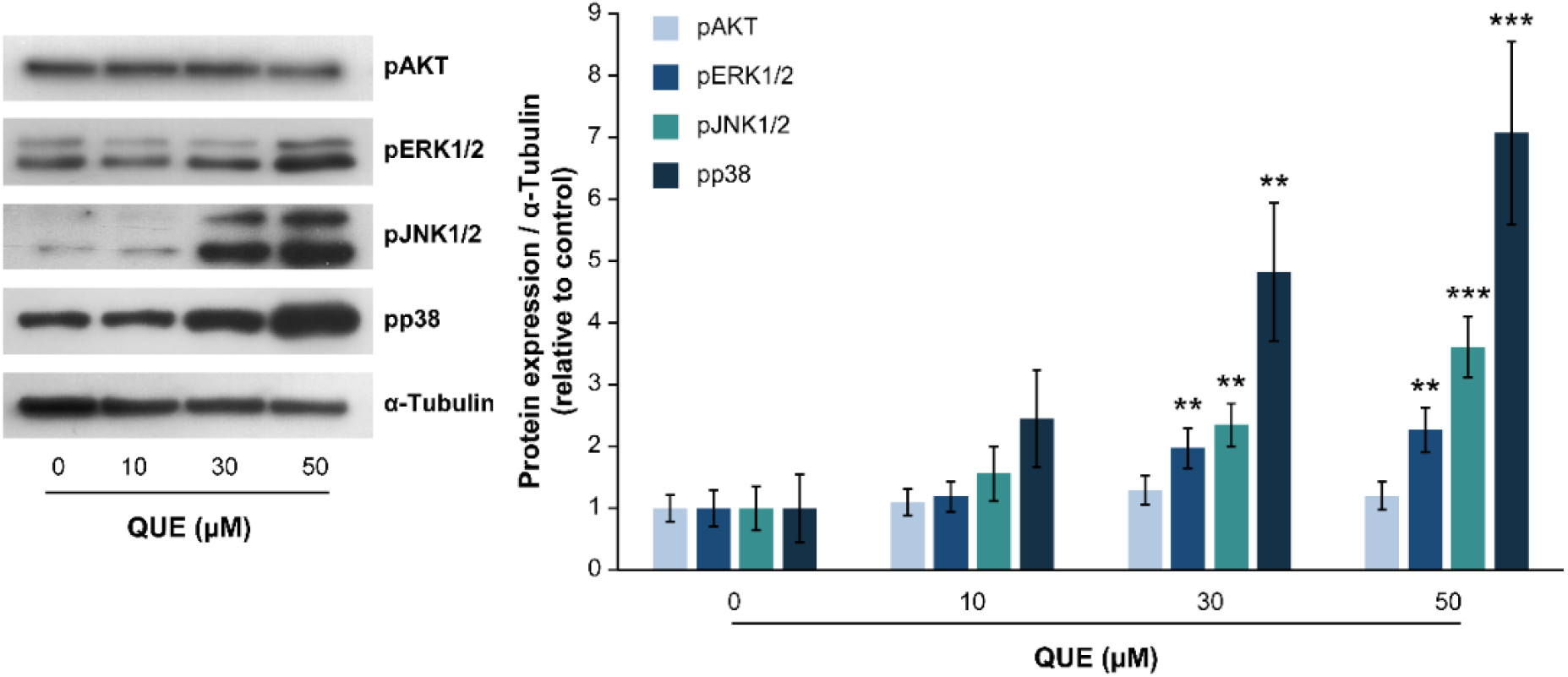
Regulation of AKT and MAPK signaling pathways by QUE. vGPCR cells were treated with QUE (10–50 µM) or vehicle (DMSO < 0.1%) for 48 h. Cell lysates were subjected to SDS-PAGE followed by Western blot analysis using antibodies against pAKT, pERK1/2, pJNK1/2, and pp38 while α-Tubulin was used as a loading control. Data from four independent experiments is expressed as percentage of the control ± SEM. Statistical analysis was performed using a linear mixed-effects model followed by Tukey’s multiple-comparisons test (**p < 0.01; ***p < 0.001).

### QUE increases oxidative stress in vGPCR cells

Although elevated ROS levels can promote carcinogenesis, they also represent a potential therapeutic target in cancer (30). In this context, QUE has been extensively studied for its ability to modulate oxidative stress and inhibit tumor progression in a concentration-dependent manner (31). Thus, it was investigated whether QUE antiproliferative action involves pro-oxidant or antioxidant activity in vGPCR cells. First, intracellular oxidant levels were assessed using the fluorogenic probe DCFDA. Cells were treated with QUE (5-45 µM) or vehicle (DMSO < 0.1%) for 24 h, while H_2_O_2_ (0.5 mM, 45 min) was used as a positive control. As shown in **Figure 3**, QUE significantly increased intracellular oxidant levels in a concentration-dependent manner.

**Figure 3.**
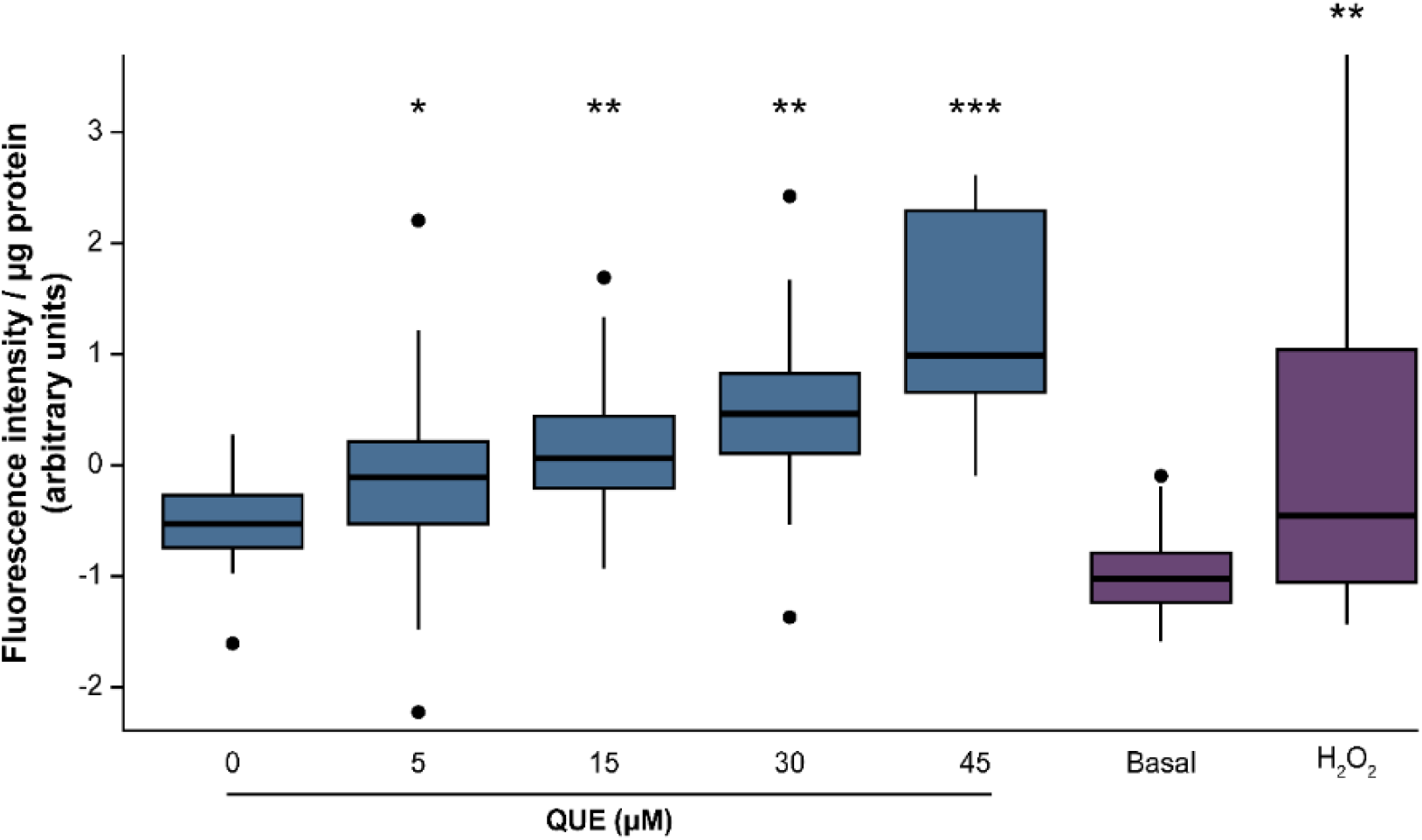
QUE induces oxidative stress in vGPCR cells. vGPCR cells were treated with QUE (5-45 µM) or vehicle (DMSO < 0.1%) for 24 h. H_2_O_2_ (0.5 mM, 45 min) was used as a positive control and compared with the basal condition (B). Intracellular oxidant levels were determined using the fluorogenic probe DCFDA. Data is presented as a boxplot from three independent experiments. Statistical analysis was performed using a linear mixed-effects model followed by Tukey’s multiple-comparisons test (*p < 0.05; **p < 0.01; ***p < 0.001).

Next, QUE’s effects on lipid peroxidation and glutathione (GSH) levels were determined using the thiobarbituric acid reactive substances (TBARS) assay and the Ellman reagent (DTNB) assay, respectively. In both cases, vGPCR cells were treated with QUE (30 µM) or vehicle (DMSO < 0.1%) for 24 or 48 h. As shown in **Figure 4A**, QUE induced a highly significant increase in lipid peroxidation after 48 h of treatment, whereas no significant differences were detected at 24 h. Similarly, QUE significantly increased total GSH levels only after 48 h of treatment (**Figure 4B**). The glutathione peroxidase (GPX) family plays a central role in detoxifying peroxides and maintaining cellular redox homeostasis (32,33). Among these enzymes, glutathione peroxidase 4 (GPX4) is a key antioxidant enzyme that reduces membrane phospholipid hydroperoxides, thereby preventing lipid peroxidation and ferroptotic cell death (33). Therefore, *Gpx4* mRNA expression was also evaluated under the same experimental conditions (**Figure 4C**). Although *Gpx4* showed a trend toward upregulation after 24 h of QUE treatment and toward downregulation after 48 h, neither change was statistically significant.

**Figure 4.**
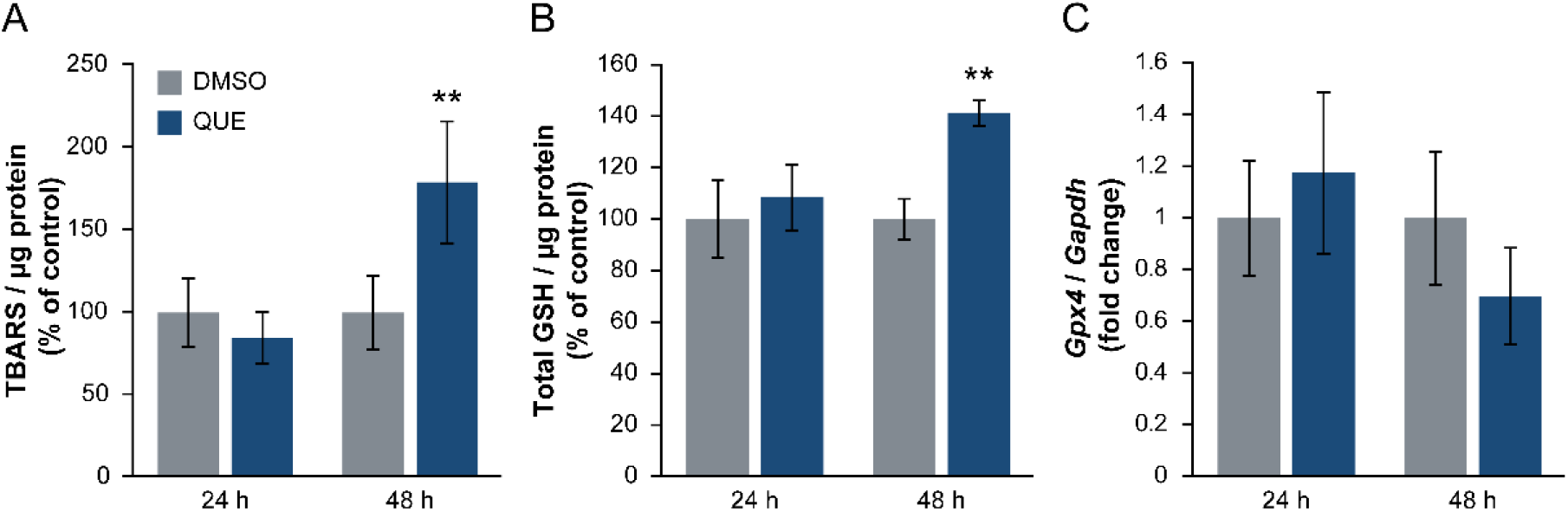
QUE promotes oxidative stress in vGPCR cells. vGPCR cells were treated with QUE (30 µM) or vehicle (DMSO < 0.1%) for 24 or 48 h. (A) Thiobarbituric acid reactive substances (TBARS) levels. (B) Total glutathione (GSH) content. (C) Gpx4 mRNA expression levels. Data represent three independent experiments. Statistical analysis was performed using the Kruskal–Wallis test (**p < 0.01).

### Role of AKT and MAPKs in QUE-induced intracellular oxidant production

Because QUE displayed a pro-oxidant effect in vGPCR cells, the potential involvement of the PI3K/AKT and MAPK pathways in this response was investigated. Cells were preincubated with specific inhibitors of PI3K/AKT (LY294002), p38 (SB203580), ERK1/2 (PD98059), or JNK1/2 (SP600125) at 10 µM for 15 min. Thereafter, QUE (30 µM) or vehicle (DMSO < 0.2%) was added, and cells were incubated for an additional 24 h without removing the inhibitors. Intracellular oxidant levels were subsequently quantified using the DCFDA assay. As shown in **Figure 5**, inhibition of either JNK1/2 or PI3K/AKT pathways significantly increased intracellular oxidant levels compared with the control condition. Furthermore, QUE treatment enhanced this effect, provoking a greater increase in oxidant levels when JNK1/2 was inhibited. A similar trend was observed following PI3K/AKT inhibition; however, this difference did not reach statistical significance.

**Figure 5.**
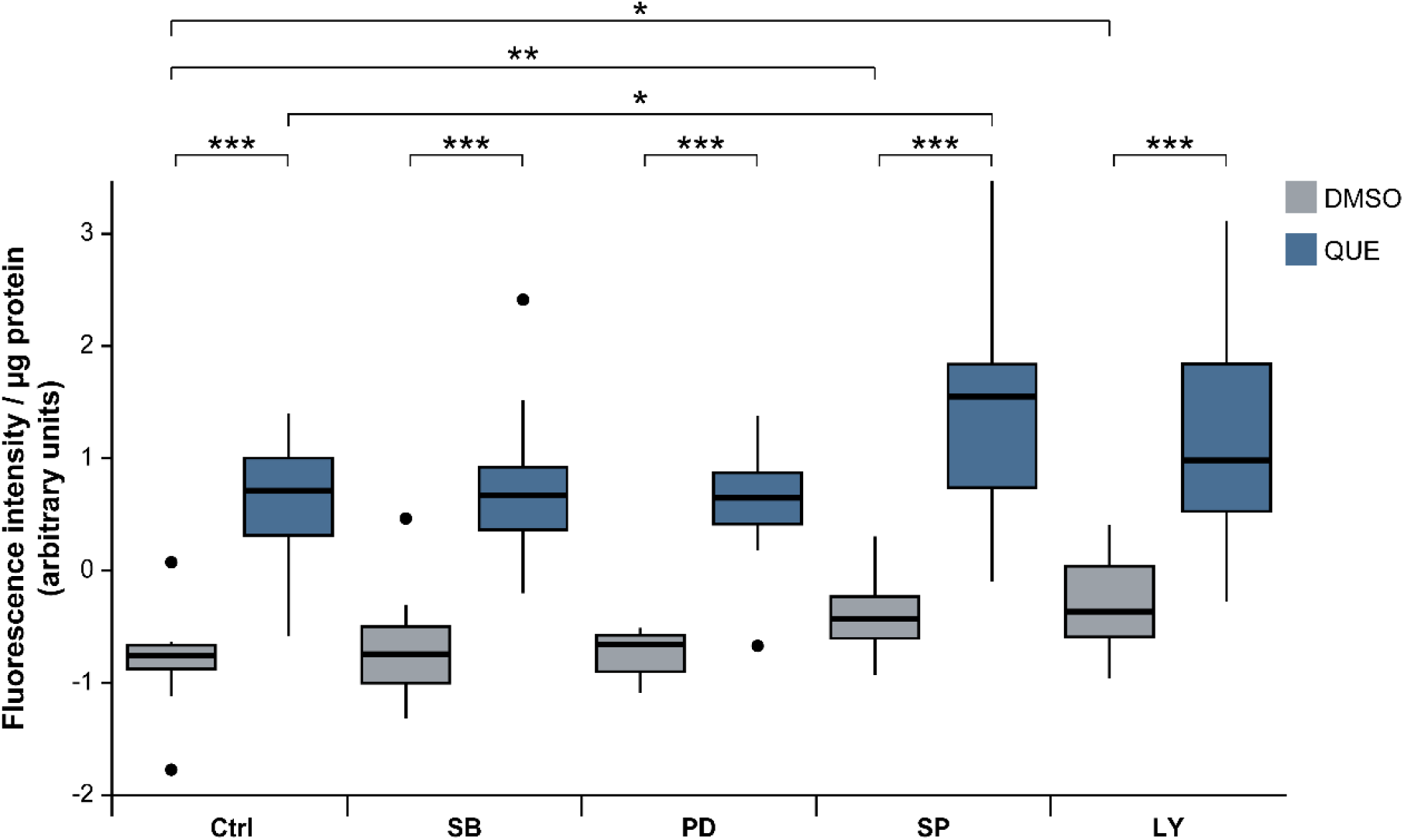
Role of AKT and MAPK pathways in QUE-induced intracellular oxidant production. vGPCR cells were treated with QUE (30 µM) or vehicle (DMSO < 0.2%) for 24 h in the presence or absence of 10 µM of inhibitors for the following kinases: p38 (SB203580), MEK/ERK (PD98059), JNK (SP600125), and PI3K (LY294002). Intracellular oxidant levels were measured using the fluorogenic probe DCFDA. Data is presented as boxplot from three independent experiments. Statistical analysis was performed using the Kruskal–Wallis and Mann–Whitney U tests (*p < 0.05; **p < 0.01; ***p < 0.001).

In parallel experiments, the phosphorylation state of AKT, p38, ERK1/2 and JNK1/2 were analyzed by Western blot. As shown in **Figure 6**, pretreatment with either SP600125 or LY294002 significantly attenuated QUE-induced phosphorylation of both JNK1/2 and ERK1/2, whereas neither inhibitor affected QUE-induced p38 activation. Similarly, PD98059 reduced the increase in ERK1/2 phosphorylation induced by QUE; however, pretreatment with SB203580 increased p38 and JNK1/2 phosphorylation, with a minor increase in ERK1/2 phosphorylation. In contrast, while neither LY294002 nor QUE alone significantly altered AKT phosphorylation, their combined treatment significantly reduced pAKT levels relative to the corresponding controls.

**Figure 6.**
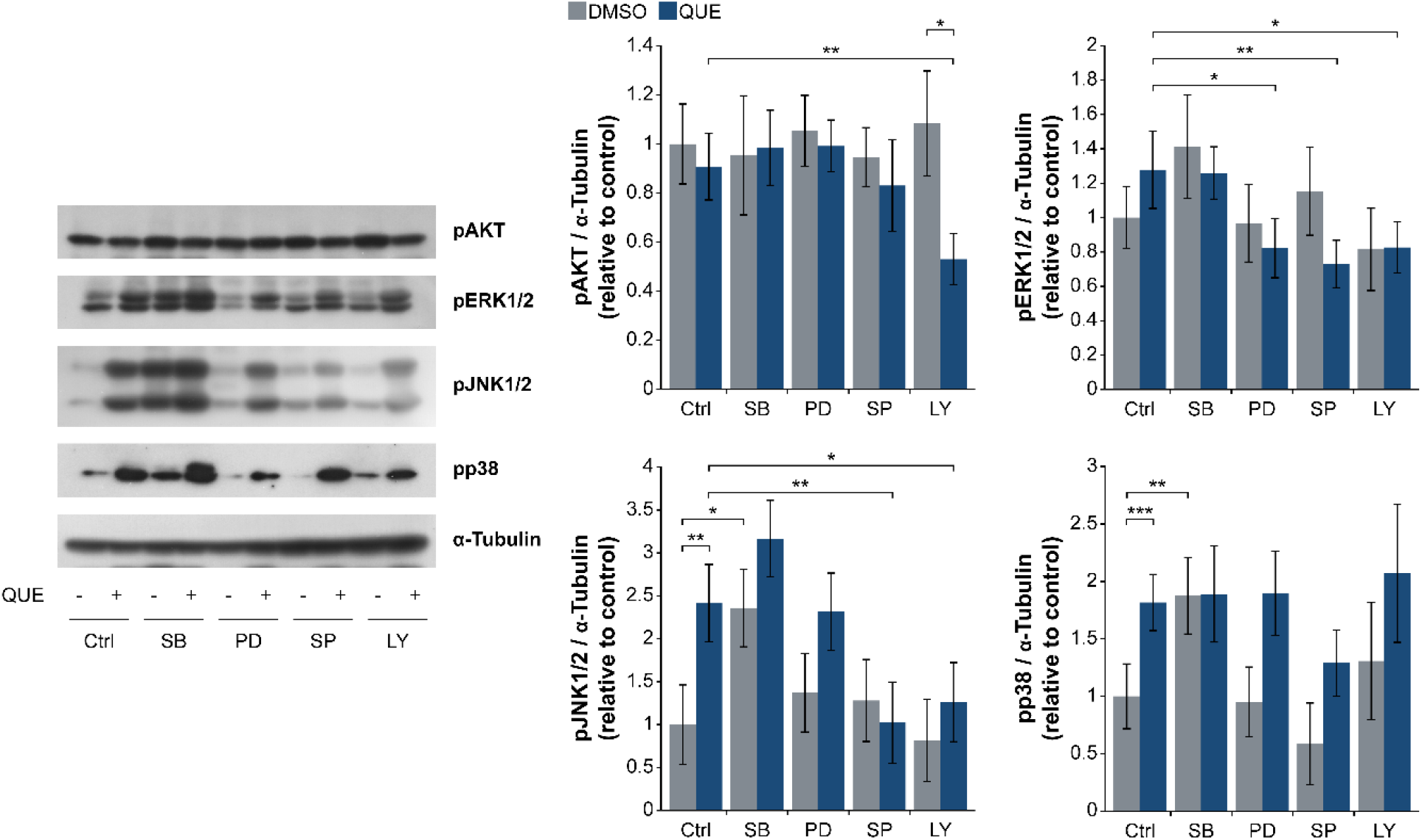
Modulation of QUE-induced MAPK activation by PI3K/AKT and MAPK inhibitors. vGPCR cells were treated with QUE (30 µM) or vehicle (DMSO < 0.2%) for 24 h in the presence or absence of 10 µM of inhibitors for the following kinases: p38 (SB203580), MEK/ERK (PD98059), JNK (SP600125), and PI3K (LY294002). Cell lysates were subjected to SDS-PAGE followed by Western blot analysis using antibodies against pAKT, pERK1/2, pJNK1/2, and pp38 while α-Tubulin was used as a loading control. Data from four independent experiments is expressed as percentage of the control ± SEM. Statistical analysis was performed using a linear mixed-effects model followed by Šídák’s multiple-comparisons test (*p < 0.05;**p < 0.01; ***p < 0.001).

### QUE induces DNA damage in vGPCR cells

Considering that QUE-induced oxidative stress may promote DNA damage (34) and taking into account previous reports describing its direct genotoxic effects (35), DNA damage was assessed using the comet assay. vGPCR cells were treated with QUE (30 µM) or vehicle (DMSO < 0.1%) for 24 or 48 h, while H_2_O_2_ (0.5 mM, 1 h) served as a positive control. DNA damage was quantified by measuring comet tail length using the OpenComet plugin for ImageJ (26). As shown in **Figure 7**, treatment with QUE for 48 h resulted in longer comet tails compared with the control condition, indicating increased DNA damage. In contrast, no significant differences were detected after 24 h of treatment.

**Figure 7.**
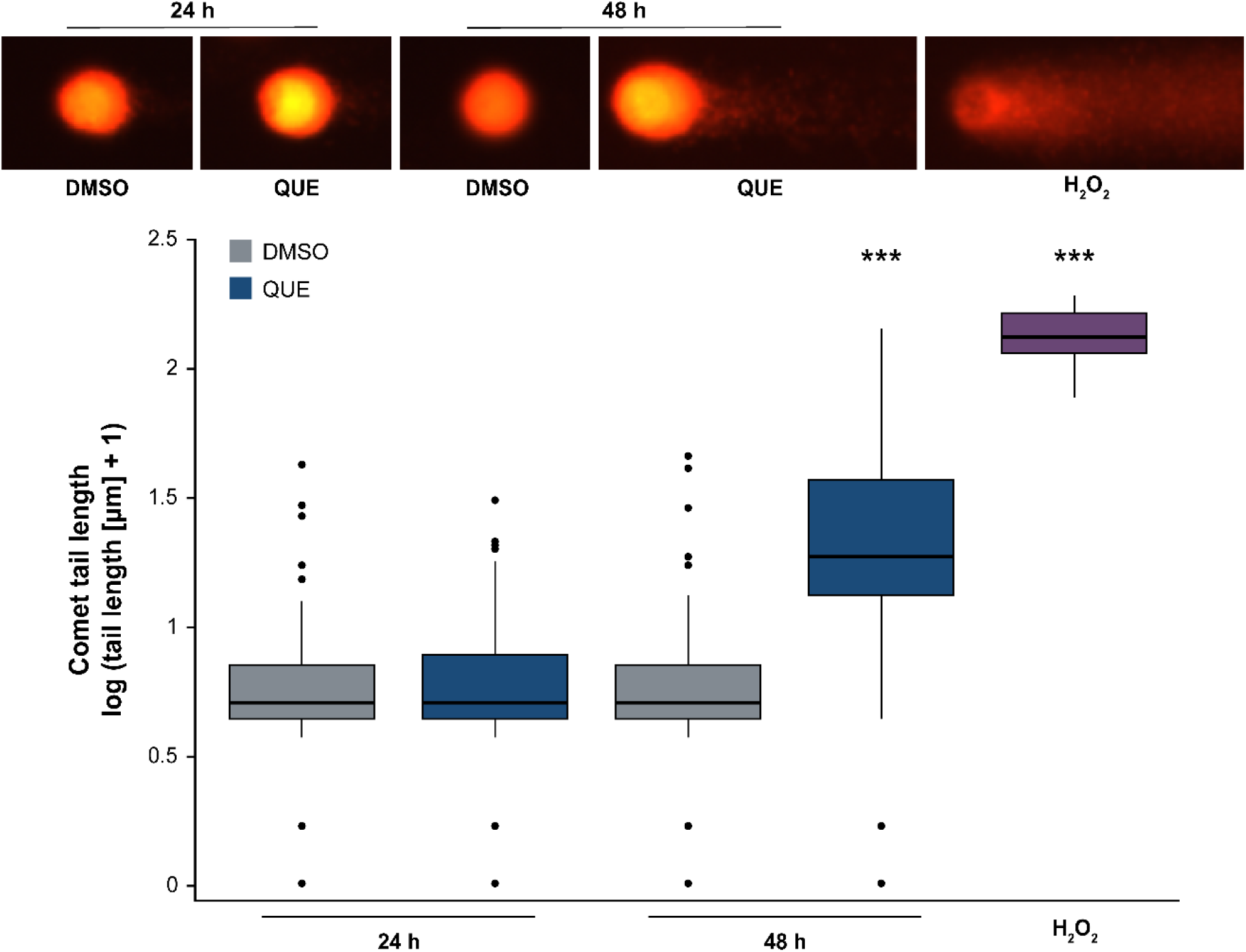
Genotoxic effect of QUE in vGPCR cells. vGPCR cells were treated with QUE (30 µM) or vehicle (DMSO < 0.1%) for 24 or 48 h. H_2_O_2_ (0.5 mM, 1 h) was used as a positive control. DNA damage was assessed using the comet assay. Fluorescence microscopy images were analyzed with the OpenComet plugin for ImageJ. Comet tail length data are presented as box-and-whisker plots. Statistical analysis of three independent experiments was performed using the Kruskal–Wallis and Mann–Whitney U tests (***p < 0.001).

## Discussion

As mentioned in the Introduction section, QUE is a dietary flavonoid with growing interest as a nutraceutical due to its multiple health-promoting properties (6). Although its antitumor activity has been extensively investigated in several cancer models, comparatively fewer studies have addressed virus-induced malignancies. Consistent with previous reports in different tumor cell lines (18,31), our group previously showed that QUE significantly reduced the proliferation and viability of vGPCR-transformed endothelial cells supporting its antineoplastic activity in this model of viral oncogenesis (20).

In this study, the effects of QUE on intracellular signaling, oxidative stress, and DNA damage in vGPCR cells were investigated. One of the most extensively studied molecular targets of QUE is the PI3K/AKT pathway. Several studies have shown that QUE suppresses PI3K/AKT signaling (36,37), an effect frequently attributed to its structural similarity to the PI3K inhibitor LY294002, which was originally developed based on the QUE scaffold (38), although both compounds interact differently with PI3K from a structural standpoint (39). Since constitutive activation of PI3Kγ and AKT is a major oncogenic mechanism downstream of vGPCR (29), inhibition of this pathway could reasonably explain the cytotoxic effects of QUE. Surprisingly, QUE did not alter AKT phosphorylation in vGPCR cells (Fig. 2), suggesting that its antiproliferative activity occurs through PI3K/AKT-independent mechanisms. Regarding the MAPK signaling pathways, previous studies have reported both inhibitory and stimulatory effects of QUE depending on the cellular context. Ki and col. (40) and Erdogan and col. (41) described inhibition of the Raf/MEK/ERK pathway, whereas other studies have shown increased ERK and JNK phosphorylation associated with apoptosis induction (42,43). The results exhibited in this work show that QUE markedly increased the phosphorylation of ERK1/2, JNK1/2, and p38 in vGPCR cells (Fig. 2), and are consistent with previous reports describing activation of MAPKs following QUE treatment (44,45), suggesting that MAPK activation may represent either an adaptive stress response or a signaling event contributing to apoptosis.

The biological activity of QUE is closely linked to its redox properties and has been reported to exhibit concentration-dependent behavior (31). At low concentrations, QUE generally exerts chemopreventive effects that have been largely attributed to its antioxidant activity. In contrast, at higher concentrations, it acts as a pro-oxidant by promoting ROS generation, thereby inducing oxidative stress and apoptosis in cancer cells (46,47). Consistent with this, our results showed that QUE significantly increased intracellular oxidant levels after 24 h of treatment (Fig. 3), whereas a significant increase in lipid peroxidation was only observed after 48 h (Fig. 4A), suggesting that oxidative damage develops progressively following the initial accumulation of ROS.

This temporal sequence supports the notion that ROS generation represents an early event in the cellular response to QUE and may act as an upstream trigger for the oxidative damage observed at later time points. Beyond their direct damaging effects, ROS are well-established signaling molecules that regulate multiple stress-responsive pathways, including MAPKs, thereby influencing cell fate decisions such as cell cycle arrest and apoptosis (48,49). Therefore, the early increase in intracellular oxidants observed in our model is likely to contribute not only to lipid peroxidation and DNA damage but also to the activation of downstream signaling events associated with the cytotoxic effects of QUE.

In parallel, QUE also increased GSH levels (Fig. 4B), a major cellular antioxidant and regulator of redox homeostasis, likely reflecting a compensatory response to the oxidative stress induced by QUE. This interpretation is consistent with previous reports describing the modulation of GSH homeostasis by QUE (50,51). Moreover, the absence of a sustained increase in *Gpx4* expression, may represent the inability of the antioxidant defense system to prevent lipid peroxidation.

Interestingly, pharmacological inhibition of either PI3K/AKT or JNK1/2 further enhanced intracellular oxidant accumulation, both in the absence and presence of QUE, indicating that these pathways partially limit oxidative stress in vGPCR cells (Fig. 5). Consistent with these findings, pretreatment with LY294002 or SP600125 prevented the QUE-induced increase in JNK1/2 and ERK1/2 phosphorylation without affecting p38 phosphorylation levels (Fig. 6). The similar effects observed after pharmacological inhibition of PI3K/AKT or JNK1/2 may be explained by functional crosstalk between these pathways, which has been previously reported in other cellular models (52,53). Thus, inhibition of either pathway may impair the cellular response to QUE-induced oxidative stress, thereby promoting intracellular ROS accumulation. On the other hand, although SB203580 and PD98059 did not significantly modify intracellular oxidant accumulation (Fig. 5), pretreatment with SB203580 increased the phosphorylation levels of p38, JNK1/2, and, to a lesser extent, ERK1/2 (Fig. 6). This effect is consistent with feedback mechanisms within the MAPKs pathways, since SB203580 inhibits the catalytic activity of p38 rather than its phosphorylation (54). Therefore, inhibition of p38 activity may lead to compensatory activation of upstream or parallel MAPK signaling, resulting in increased phosphorylation of p38, JNK1/2, and ERK1/2. In addition, PD98059 reduced ERK1/2 phosphorylation in the presence of QUE without affecting JNK1/2 phosphorylation levels (Fig. 6). Taken together, these findings suggest that the cellular response to QUE-induced oxidative stress in vGPCR cells relies mainly on JNK1/2 activity, with a possible secondary contribution of ERK1/2 that is insufficient to counteract the cytotoxic effects of QUE at the tested concentration.

Finally, the increased oxidative stress induced by QUE was accompanied by significant DNA damage after 48 h of treatment (Fig. 7), supporting the hypothesis that ROS generation promotes genomic damage and may contribute to the subsequent induction of apoptosis, as previously reported in other cancer models (55).

## Conclusion

In conclusion, QUE inhibits the proliferation of vGPCR-transformed endothelial cells through a mechanism involving oxidative stress. The increase in oxidant levels was associated with MAPK activation, particularly JNK1/2. Furthermore, the imbalance between oxidant production and antioxidant defenses was accompanied by DNA damage, suggesting that oxidative stress contributes to the antiproliferative and proapoptotic effects of QUE in this model.

## Abbreviations

AIDS: acquired immunodeficiency syndrome
AKT: protein kinase B
DCFDA: 2′,7′-dichlorofluorescein diacetate
DMSO: dimethyl sulfoxide
DTNB: 5,5′-dithiobis(2-nitrobenzoic acid)
ERK1/2: extracellular signal-regulated kinases 1/2
GSH: glutathione
HHV-8: human herpesvirus 8
HIV: human immunodeficiency virus
IC_50_: half maximal inhibitory concentration
JNK: c-Jun N-terminal kinase
KS: Kaposi’s sarcoma
KSHV: Kaposi’s sarcoma-associated herpesvirus
MAPK: mitogen-activated protein kinase
PI3K: phosphoinositide 3-kinase
QUE: quercetin
ROS: reactive oxygen species
SV40: simian virus 40
TBARS: thiobarbituric acid reactive substances
vGPCR: viral G protein-coupled receptor.

