## Supplementary Material for "Quercetin alters redox homeostasis, induces MAPK activation and promotes DNA damage in vGPCR-transformed endothelial cells"

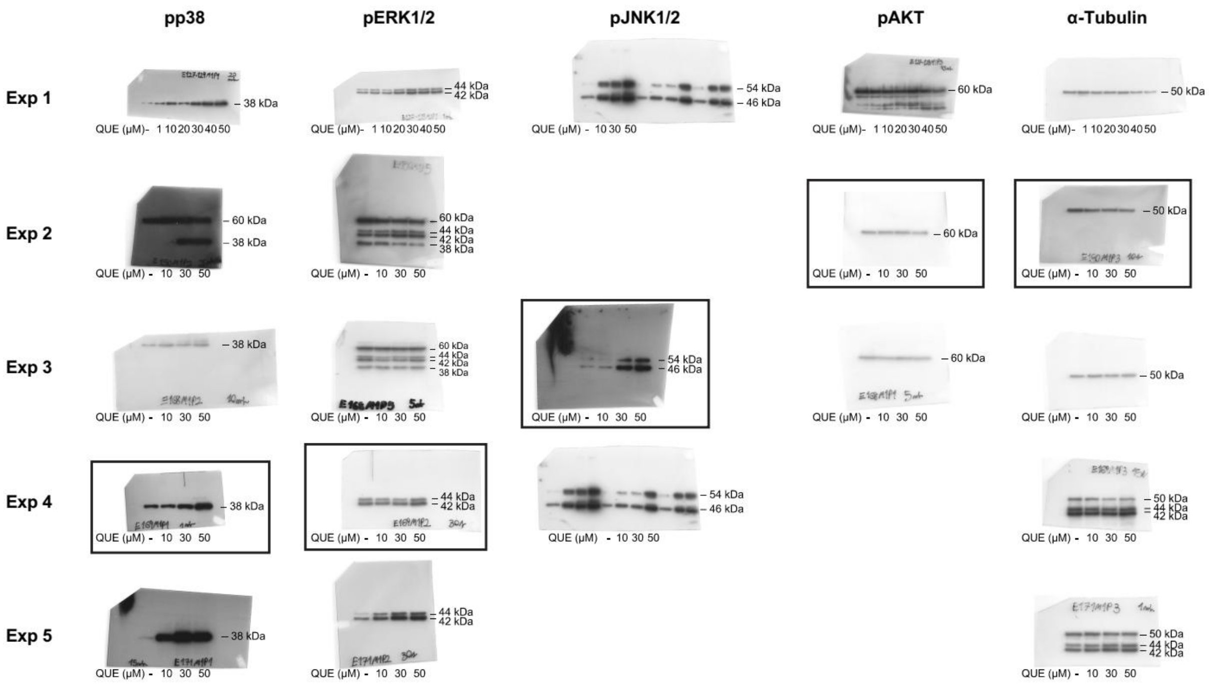


**Supplementary Figure S1. Independent Western blot replicates corresponding to Figure 2: Regulation of AKT and MAPK signaling pathways by QUE.** Black boxes indicate the blots used for the corresponding figure.

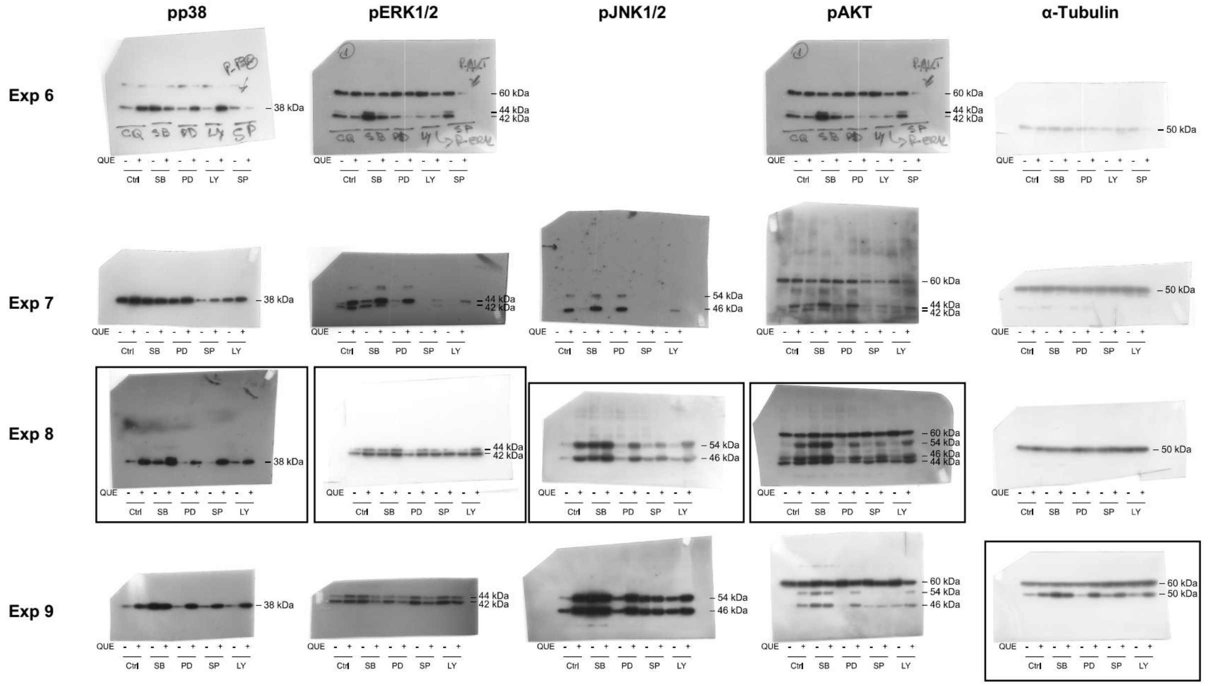


**Supplementary Figure S2. Independent Western blot replicates corresponding to Figure 6: Modulation of QUE-induced MAPK activation by PI3K/AKT and MAPK inhibitors.** Black boxes indicate the blots used for the corresponding figure.
